# NRSF–mediated repression of neuronal genes in developing brain persists in the absence of NRSF-Sin3 interaction

**DOI:** 10.1101/245993

**Authors:** Alicia M. Hall, Annabel K. Short, Akanksha Singh-Taylor, Jennifer Daglian, Tadashi Mishina, William K. Schmidt, Hiroyuki Kouji, Tallie Z. Baram

**Author notes:** These authors contributed equally to this work (Alicia M. Hall, Annabel K. Short).

## Abstract

Repression of target genes by the transcriptional repressor neuronal restrictive silencing factor (NRSF)/repressor element 1 silencing transcription factor (REST) contributes to enduring plasticity in the developing brain. However, the cofactor(s) interacting with NRSF to enable target gene repressor are not well understood, and may vary among neuronal populations and brain regions as well as with different contexts. Here we employed the novel designer drug mS-11 to block the interactions of the cofactor Sin3 with NRSF. We tested if NRSF-Sin3 interaction is required for repression of NRSF target genes in developing hypothalamus after activity-dependent modulation of NRSF function. In the hypothalamus *in vitro*, blocking glutamatergic neurotransmission robustly increased NRSF binding to the target gene *Crh*, resulting in its repression. Blocking the binding of NRSF to the chromatin with decoy NRSE-oligodeoxynucleotides abrogated this repression. In contrast, mS-11 at several concentrations did not impede *Crh* repression. NRSF-mediated repression may underlie disease processes such as the onset of epilepsy. Therefore, identifying small-molecule antagonists of NRSF is crucial for the development of disease-preventing or modifying interventions.

## Introduction

The transcriptional repressor neuronal restrictive silencing factor (NRSF or REST) is emerging as an important mediator of neuronal plasticity (1–3). NRSF expression was originally described in non-neuronal tissues where it suppresses neuron-specific genes (4–6), indicating that many neuronal genes must carry NRSF-response elements (NRSE/RE1) and are repressed by augmented NRSF levels or activity (3,6). A second important role for NRSF was established by showing that transition of neuronal precursors to mature neurons requires dissociation of NRSF from the chromatin of neuronal genes (7). More recently, NRSF expression in mature neurons has been described, where the factor may be crucial for normal function (8–10). In differentiated neurons, NRSF is especially crucial to those still developing (11), where expression of NRSF-regulated genes contributes to several aspects of maturation, including development of excitatory synapses (1,2,12,13). This is important because many neuropsychiatric disorders seem to originate during the developmental epoch (infancy and early childhood) when neurons are involved in the creation of synaptic contacts and their refinement and pruning to mature circuits (12,14–17).

Indeed, recent evidence suggests that blocking the binding of the transcription factor NRSF to the chromatin in rodent models, ameliorates the development of epileptic seizures, and prevents the development of memory problems, which are seen after insults in humans (18–20). In experimental models, NRSF function was blocked using the administration of decoy oligodeoxynucleotide (ODNs) directly into the brain. However, whereas blocking NRSF holds great promise to modulate epileptogenesis in humans, the administration of ODNs is not clinically practical. These facts provide great impetus for the identification and/or synthesis of small-molecule ‘designer drugs’ to block NRSF actions.

NRSF exerts its repressive effects on gene expression by interacting with RE1/NRSE sites and recruiting corepressors including Sin3A,B (Sin3) (7,21–23) G9a and/or CoREST (24–26). These proteins then engage additional transcriptional regulators and cofactors including the G9a histone methyltransferase (27), the histone deacetylases 1 and 2, and the H3 lysine 4 demethylase LSD1 (28), as well as others (29). The interacting cofactors engage distinct domains of the NRSF molecule (Fig 1). Our initial work suggested that cofactors targeting the N-terminus of NRSF might contribute importantly to the actions of the repressor in the postnatal developing brain (18,30). Therefore, we tested the potential role of Sin3, an NRSF-N-terminus interacting cofactor (21), in contributing to NRSF-mediated gene repression in the developing brain. We capitalized on the well-established function of NRSF in the repression of one of its target genes, corticotropin releasing factor, encoded by *Crh* (31,32) in the hypothalamus. Thus, we employed an *in vitro* hypothalamic slice-culture system that maintains the integrity of hypothalamic three-dimensional structure yet provides a controllable system. We tested if blocking NRSF-Sin3 interaction, using the small molecule mS-11, prevents the repression of CRH expression that is observed upon reduction of glutamatergic neurotransmission. In addition, we compared the efficacy of mS-11 to that of an NRSE-ODN that directly interferes with binding of NRSF to the chromatin.

**Fig 1.**
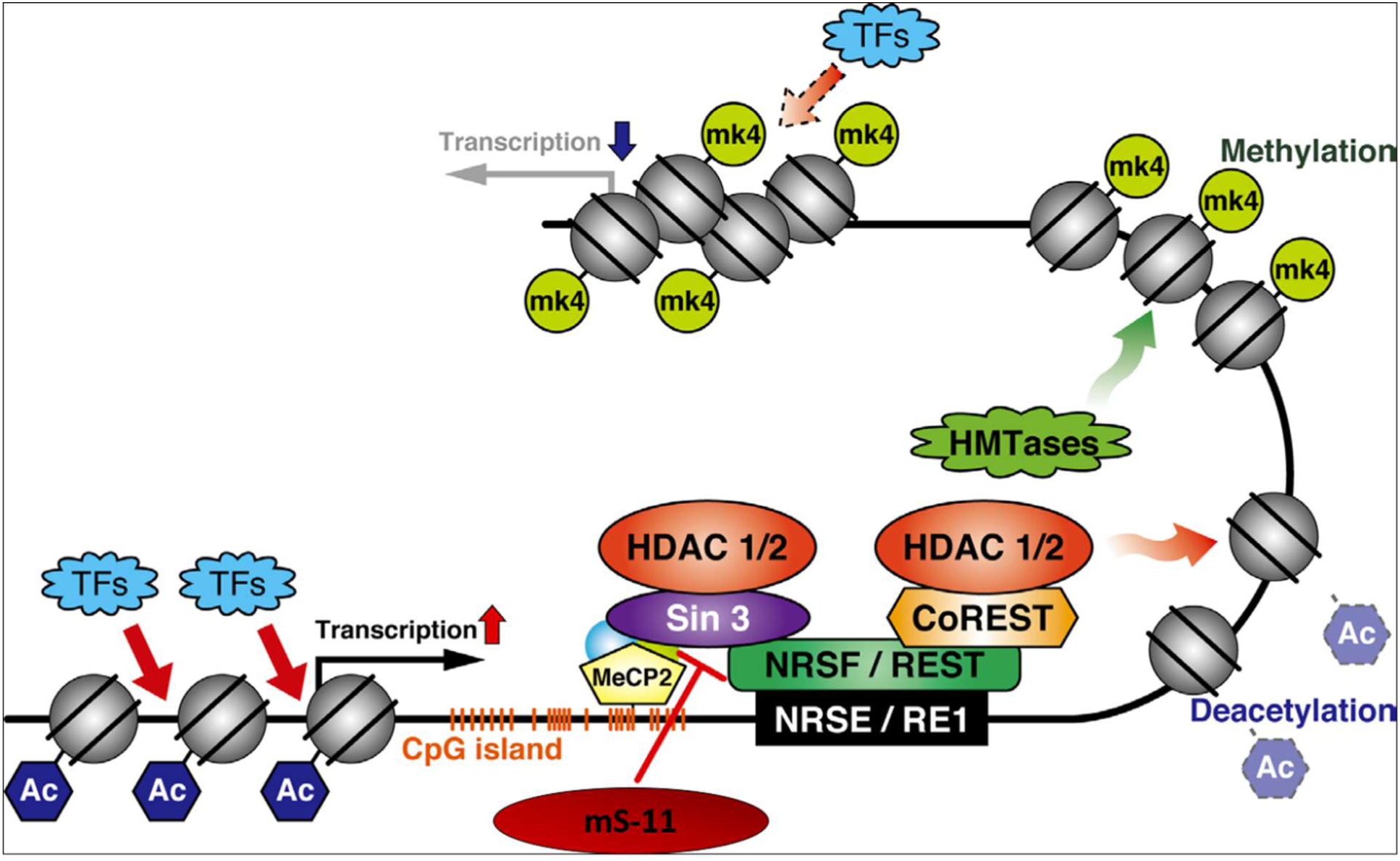
NRSF binds directly to the DNA. It recruits cofactors including Sin3 to its N-terminus domain and CoREST to its C-terminus domain. These complexes are thought to enable NRSF regulation of gene expression.

## Materials and Methods

### Drug development

Computer-driven 3D methodologies were used to design a molecule that mimics the NRSF N-terminus moiety that interacts with the mammalian Sin3B-PAH1 domain. An improved compound called mS-11 was identified. Further details of the compound and its synthesis and testing are provided in Ueda et al., 2017. Because the homology between Sin3A and Sin3B is quite high in the PAH1 domain, mS-11 is expected to interfere with NRSF binding to Sin3A as well.

### *In vitro* system for NRSF-mediated gene repression

Organotypic hypothalamic slice cultures were generated from rats as described previously (8). Animals were handled according to NIH guidelines for care and use of laboratory animals and in accordance with protocol approval from the University of California-Irvine Institutional Animal Care and Use Committee. Subjects were progeny of timed-pregnant Sprague-Dawley rats. Rats were housed under a 12-hour light-dark cycle in humidity and temperature controlled rooms, with *ad libitum* access to food and drinking water. Parturition was checked daily, and the day of birth was considered postnatal day (P) 0. To generate of hypothalamic explant cultures, pups were sacrificed on P6-7.

Cultures were prepared using a stationary hypothalamic slice culture protocol as previously described (8). Rat pups were decapitated on P6-P7 (weight range 13-17g) (day in vitro (DIV) 0), brains were removed, and hypothalamic blocks were dissected and cut into 350 μm coronal sections on McIlwain tissue chopper in ice cold prep media (MEM (Thermo Fisher Scientific, Waltham, MA)), L-Glutamine (Thermo Fisher Scientific, Waltham, MA), HEPES Buffer (Thermo Fisher Scientific, Waltham, MA), glucose, and cell culture grade water (GE Healthcare, Little Chalfont, UK) in a laminar flow hood. Sections containing the PVN were then maintained on 0.4 µm, 30 mm diameter cell culture inserts (Merck Millipore, Darmstadt, Germany) in six well plates with culture media (Minimum Essential Medium, Hank’s balanced salt solution, heat-inactivated horse serum, HEPES, glucose, glutamine, ascorbic acid, insulin, NaHCO3, and cell culture grade water). Explants were maintained for 8 days at 37 °C in 5% CO2 enriched air in an incubator, with the media refreshed every 48 hours (h). Explants were kept in serum-containing medium from DIV 0 to DIV 6, which also contained antibiotics (penicillin and streptomycin (Thermo Fisher Scientific, Waltham, MA) until DIV 4. On DIV 6 cultures were transferred to serum free media (Minimum Essential Media, HEPES, glucose, glutamine, ascorbic acid, insulin, NaHCO3, cell culture grade water), and treated with either sterile nuclease-free tissue-culture grade water (vehicle), or 50 µM MK-801 (Sigma-Aldrich, St. Louis, MO) and 50 µM CNQX (Sigma-Aldrich, St. Louis, MO). Media containing either vehicle or glutamate receptor antagonists (GluRs) were refreshed every 12 h for 52 h. After 52 h, cultures were then harvested on dry ice, and stored at - 80°C until further processing.

### Blocking of NRSF function using NRSE-ODNs

For oligodeoxynucleotide (ODN) treatment, previously described phosphothiolated ODNs consisting of either a control, randomly ordered sequence (scrambled; SCR) or a sequence coding for NRSF binding site (NRSE), were used (3,18). NRSE-ODN sequence: 5’-GGAGCTGTCCACAGTTCTGAA-3’ and scrambled ODN sequence: 5’-AGGTCGTACGTTAATCGTCGC-3’ (Sigma). Slices were maintained as described above, transferred to serum free medium on DIV 6, and exposed to one of the following treatments: [Veh + NO ODN], [Veh + SCR], [Veh + NRSE], [CNQX/MK-801+NO ODN], [CNQX/MK-801 + SCR], or [CNQX/MK-801 + NRSE]. Dose-response analysis determined 10 nM as the optimal ODN concentration. The ODNs were used only for the first 12 h of treatment. Media containing either vehicle or antagonists were refreshed every 12 h for 52 h. Cultures were then harvested and stored as described above. We have previously shown that the ODNs enter neuronal cytoplasm and nuclei (8,20).

### Blocking of Sin3-NRSF interaction using mS-11

The protocol for using mS-11 followed the one described above. Briefly, hypothalamic explants were kept in culture from DIV 0 to DIV 6. On DIV 6 cultures were treated with either sterile nuclease-free tissue-culture grade water (vehicle), or 50 µM MK-801 and CNQX to block ionotropic glutamate receptors. Media containing either vehicle or glutamate receptor (GluR) antagonists were refreshed every 12 h for 52 h. The experimental groups included: [Veh + diluent], [Veh + mS-11], [CNQX/MK-801 + diluent] or [CNQX/MK-801 + mS-11]. Two concentrations of mS-11 were used, and mS-11 was diluted in 0.1%DMSO and applied directly to the medium for final concentrations of 10 µM or 70 µM. The latter aimed to approximate concentrations achieved upon *in vivo* administration of the drug, which was given at 30 mg/kg (PRISM Co. Material, unpublished). Media containing either vehicle, antagonists, mS-11 or antagonists and mS-11, were refreshed every 12 h for 52 h. Cultures were then harvested and stored as described above.

### RNA extraction, reverse transcription, and qRT-PCR

Samples were thawed on ice, and RNA was isolated using RNeasy Mini Kit (Qiagen, Valencia, CA) as per manufacturer’s protocol. RNA was quantified, and purity was analyzed using NanoDrop (Thermo Fisher Scientific, Waltham, MA). It was immediately converted to cDNA with random hexamers using first strand cDNA synthesis kit (Roche, Basel, Switzerland) following manufacturer’s protocol. qRT-PCR was performed using SYBR Green chemistry (Roche, Basel, Switzerland) on a Lightcycler 96 (Roche, Basel, Switzerland) with primers for specific transcripts. Gapdh or *β-Actin* served as the internal control (‘housekeeping gene’), and relative quantification of mRNA expression was accomplished using the cycle threshold method (2ˆ-ΔΔCt). Minus-reverse transcription and non-template controls were routinely used to eliminate the possibility of genomic contamination or false positive analyses. Primer sequences used for qRT-PCR have been published (8).

### Statistical considerations and analyses

All analyses were conducted without knowledge of treatment group. Analysis of variance and Tukey’s multiple comparison post-hoc tests were employed as required using GraphPad PRISM software.

## Results

### NRSF mediates the repression of its target gene, *Crh*, that is provoked by reduction of glutamatergic neurotransmission

In accord with our prior observations (Korosi et al., 2010; A Singh-Taylor et al., 2017), blocking of excitatory neurotransmission using a cocktail of glutamate receptor (GluR) antagonists (CNQX/MK-801) to CRH–expressing cells in hypothalamic cultures reduced Crh mRNA levels by around 50% (Fig 2). Interfering with the binding of NRSF to its recognition site on the chromatin using NRSE-ODN decoys (Fig 2A) abrogated this repression of the *Crh* gene (Fig 2B; Significant interaction on two-way ANOVA; F[2,77]= 7.09, p<0.01; post hoc, *p<0.05). This was specific to NRSF, because a random-sequence (SCR)-ODN had negligible effect on the suppression of *Crh* expression (Figs 2B).

**Fig 2.**
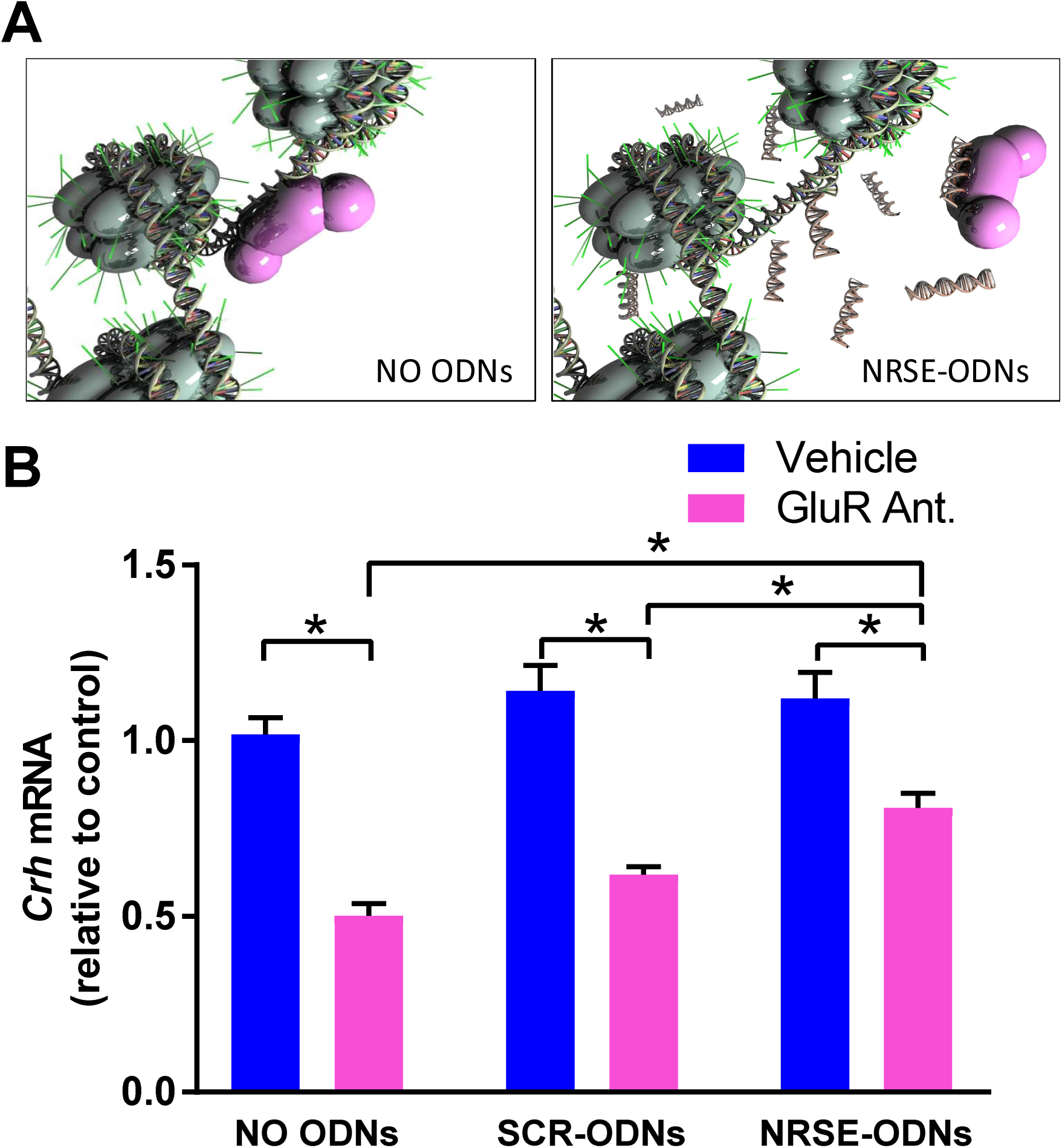
Blocking NRSF binding to the chromatin using NRSE-ODNs prevents the GluR antagonist–induced repression of the *Crh* gene. (A) NRSE-ODNs act as decoys: they bind NRSF and thus prevent it from binding to RE1/NRSE element on the gene. (B) qPCR for *Crh* mRNA levels. RNA isolated from hypothalamic cultures after 52 hours and normalized to *Gapdh*. GluR antagonists (CNQX/MK-801) reduced *Crh* expression levels and NRSE-ODNs prevented this reduction. This was not evident after application of random (scrambled, SCR) ODNs. Two-way ANOVA; significant interaction, F_[2,77]_= 7.09, p<0.01; post hoc, *p<0.05 ^#^p<0.09; n=11- 21/group. *A) Used under Creative Commons license attributed to Shawn McClelland*

These finding indicated that *Crh* mRNA repression in developing hypothalamus by blocking ionotropic glutamate receptors is a robust and reliable system, which recapitulates an *in vivo* repression by early-life experience (33,35–37). Further, the *Crh* mRNA repression requires the function of NRSF. We then set out to determine if the interaction of NRSF with the cofactor Sin3 is required for NRSF to repress *Crh*. This was accomplished by the use of a compound designed to mimic the Sin3B-PAH1–interacting domain of NRSF (38).

### mS-11 does not prevent repression of *Crh* mRNA after ionotropic glutamate receptor blockade

To examine if Sin3-NRSF interaction is required for the NRSF-mediated repression of Crh mRNA expression in hypothalamus, we incubated hypothalamic explants with two distinct concentration of the mS-11. As is shown in Fig 3, blocking ionotropic glutamate receptors (using CNQX/MK-801) robustly repressed CRH mRNA levels. However, this reduction was not influenced by either 10 µM (Fig 3A; Significant main effect of GluR antagonists on two-way ANOVA; F[1,12]= 104.5, p<0.001) or 70 µM concentrations of mS-11 (Fig 3B; Significant main effect of GluR antagonist on two-way ANOVA; F[1,12]= 15.93, p<0.01). The higher concentration is well in line with those employed in vivo (38) and in vitro (PRISM Co. Datasheet, H. Kouji, personal communication). In addition, this concentration resulted in apparent toxicity within the cultures, suggesting that it is limiting.

**Fig 3.**
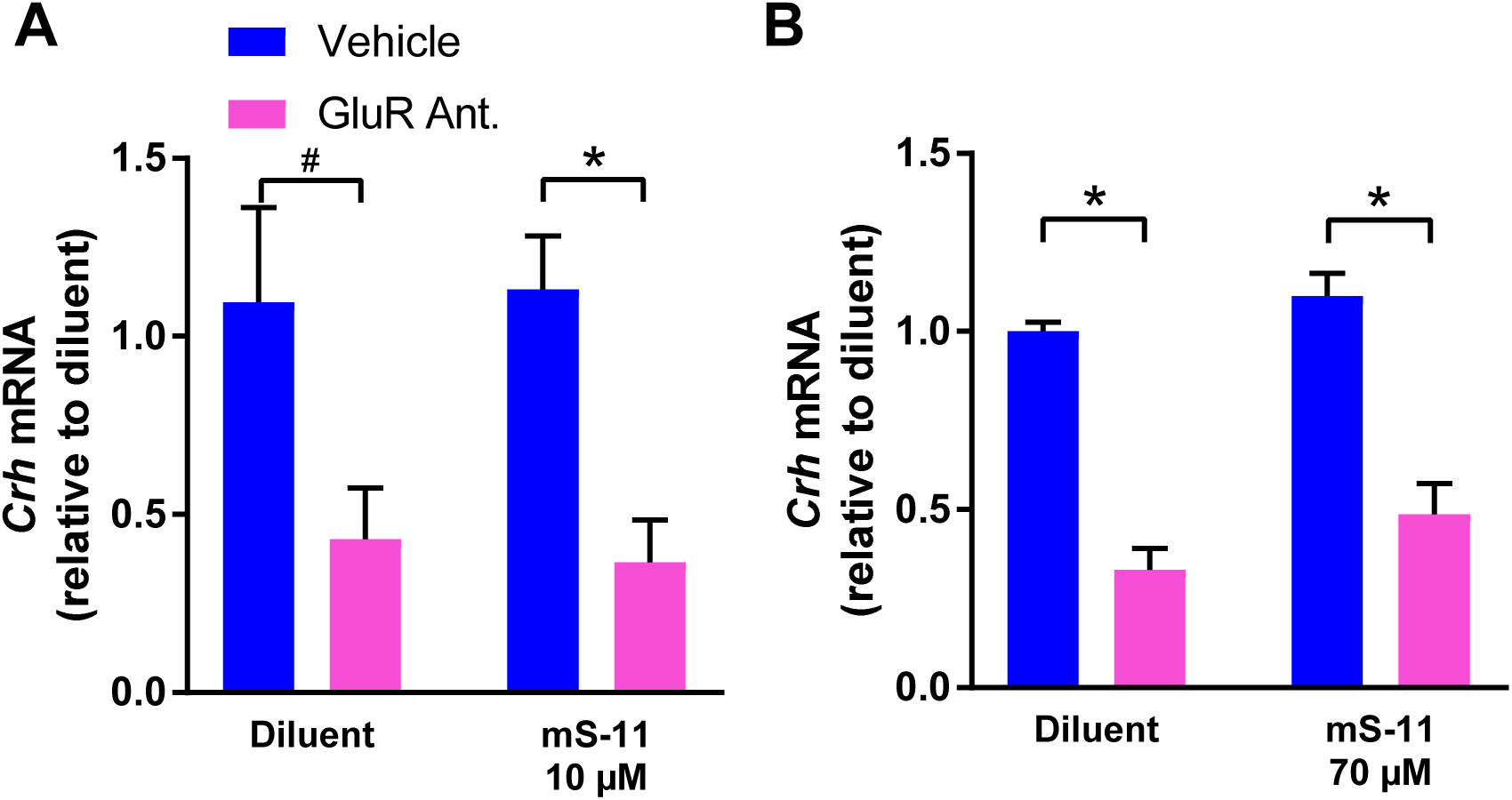
mS-11 did not block GluR antagonist–induced repression of the *Crh* gene. Shown are results obtained using qPCR for *Crh* mRNA levels: RNA was isolated from hypothalamic cultures at 52 hours and was normalized to *β-actin*. (A) GluR antagonists (CNQX/MK-801) reduced *Crh* expression levels and mS-11 had no effect on this reduction at either 10 µM (two-way ANOVA; significant main effect of GluR antagonists F[1,12]= 104.5, p<0.001; n=4/group). (B) or 70 µM (two-way ANOVA; significant main effect of GluR antagonists, F_[1,12]_= 15.93, p<0.01; n=4/group).

## Discussion

The principal findings in these series of experiments are (1) NRSF binding to cognate recognition sites within the *Crh* gene contributes critically to NRSF-mediated repression of this target gene in the developing brain; (2) interaction and recruitment of the N-terminus cofactor Sin3 may not be necessary for this repression.

The mechanisms by which experience, environment and activity early-in life lead to persistent, epigenetic changes in gene expression are a topic of major importance (32,39,40). These epigenetic changes may be critical components of neurodevelopmental as well as later-onset brain disorders (15,17,41,42). Networks of transcription factors and cofactors, together with chromatin modulations (7,24,25,43) and numerous other signals mediate the enduring effects of early-life experiences on gene expression in specific neurons (30,44).

Whereas the involvement of NRSF in neuronal plasticity in developing hippocampus (20,26,45) and hypothalamus (8) is established, the molecular machinery engaged by NRSF is not well understood. NRSF recruits a number of molecules and molecular complexes (22,25,26,29), which may contribute to initiation of gene repression. For some genes, the level of expression is adjusted further in neurons by CoREST/MeCP2 repressor complexes as well as others that remain bound to a site of methylated DNA distinct from the RE1 site (7,18). Therefore, direct interference with NRSF-binding may not be specific to a given gene set.

To determine therapeutic approaches of NRSF interference that are applicable to clinical use, it is important to determine the mechanisms by which NRSF interacts with cofactors to repress gene expression. Current manipulations of NRSF-binding in animal studies involve administration of genetic material directly to the brain (3,18), however these are not appropriate for clinical translation. Sin3 is an attractive candidate for interfering with NRSF function. The N-terminal repressor domain of NRSF recruits this corepressor (also called mSin) which comes in two isoforms (A and B). Sin3A is recruited by NRSF in hippocampus (22). In addition, Sin3 is involved in the interaction of NRSF and the huntingtin gene product (46). This co-repressor consists of four paired amphipathic helix (PAH) domains called PAH1-PAH4. The structure of the PAH1 domain of mSin3B in complex with the N-terminal repressor of NRSF has been resolved, providing the template for the synthesis of mS-11. Ueda et al., studied the chemical shift perturbations (CSPs) of the PAH1 domain of Sin3 alone or bound to either NRSF of mS-11. The results strongly supported the notion that mS-11 would supplant NRSF-binding to Sin3, thus interfering with NRSF-Sin3 interaction (38).

In the current experiments, we employed mS-11 at high concentrations, which are larger than those tested by Ueda, and should lead to complete displacement of NRSF from the interacting PAH1 domain of Sin3. Therefore, the current experiments suggest that Sin3 is not required for the actions of NRSF in Crh-repression in developing hypothalamus. Thus, whereas the results of these experiments are limited lack of definitive evidence that mS-11 displaced NRSF from Sin3 in our system, the high concentrations we employed and the absence of any evidence of dose-response, together with Ueda et al.’s finding strongly suggest that mS-11 displaced NRSF from Sin3 in the organotypic slice culture. In addition, we employed mS-11 in a concentration that began to exert toxicity on brain tissue. Thus, aiming to improve disruption of Sin3-NRSF interaction using higher mS-11 concentration is not practical.

In conclusion, repression of target genes by NRSF contributes to enduring plasticity in the developing brain. Because NRSF-mediated repression may also underlie disease processes such as the onset of epilepsy, identifying small-molecule antagonists of NRSF is crucial for the development of disease-preventing or modifying interventions. Here we tested if interfering by the interactions of NRSF with its cofactor Sin3 might provide a therapeutic avenue, using a novel computer-designed small molecule drug. Whereas our data do not support a role of Sin3-NRSF interactions in repression of NRSF target genes in developing hypothalamus, our approach sets the stage for moderate-throughput future drug screening.

## Acknowledgements

We thank Gissell Sanchez for excellent technical help. Research supported by NIH grants NS35439, NS78279.

